# Characterisation of the UK honey bee (*Apis mellifera*) metagenome

**DOI:** 10.1101/293647

**Authors:** Tim Regan, Mark W. Barnett, Dominik R. Laetsch, Stephen J. Bush, David Wragg, Giles E. Budge, Fiona Highet, Benjamin Dainat, Joachim R. de Miranda, Mark Blaxter, Tom C Freeman

## Abstract

The European honey bee (*Apis mellifera*) plays a major role in pollination and food production, but is under threat from emerging pathogens and agro-environmental insults. As with other organisms, honey bee health is a complex product of environment, host genetics and associated microbes (commensal, opportunistic and pathogenic). Improved understanding of bee genetics and their molecular ecology can help manage modern challenges to bee health and production. Sampling bee and cobiont genomes, we characterised the metagenome of 19 honey bee colonies across Britain. Low heterozygosity was observed in bees from many Scottish colonies, sharing high similarity to the native dark bee, *A. mellifera mellifera*. Apiaries exhibited high diversity in the composition and relative abundance of individual microbiome taxa. Most non-bee sequences derived from known honey bee commensal bacteria or known pathogens, e.g. *Lotmaria passim* (Trypanosomatidae), and *Nosema spp*. (Microsporidia). However, DNA was also detected from numerous additional bacterial, plant (food source), protozoan and metazoan organisms. To classify sequences from cobionts lacking genomic information, we developed a novel network analysis approach clustering orphan contigs, allowing the identification of a pathogenic gregarine. Our analyses demonstrate the power of high-throughput, directed metagenomics in agroecosystems identifying potential threats to honey bees present in their microbiota.

---

The European honey bee, *Apis mellifera* Linnaeus, has a global distribution and a major role in pollination and food production (1). Like other pollinators, honey bee populations face multiple threats. The UN’s food and agriculture organisation estimate that 75% of all pollinators are in decline globally and their numbers have dropped by about one third in the past decade (2). Whilst flowering crops benefit greatly from a diversity of insect pollinators (3), managed honey bees are a major global contributor, providing nearly half of the service to all insect-pollinated crops on Earth (4, 5). Despite the recent increase in non-commercial beekeeping, the number of managed honey bee colonies is growing more slowly than agricultural demand for pollination (6). The decline in pollinators is not thought to be caused by a single factor but may be driven by a combination of habitat fragmentation, agricultural intensification, pesticide residue accumulation, new honey bee pests and diseases, and suboptimal beekeeping practices (7–9). Trade in honey bees from different regions of the globe have unquestionably contributed to a rise in infectious disease and there may be transmission between honey bees and wild pollinators (10, 11).

In the UK the genetic structure of honey bee populations has undergone large changes over the last 100 years. The native M-lineage subspecies, *A. m. mellifera*, had predominated in the UK, but the population was decimated in the early 20th century by a combination of poor weather and chronic bee paralysis virus, thought to have been caused by Isle of Wight disease (12). Following this, the practice of bee importation increased dramatically. In the UK today there is a growing industry that imports bees from mainland Europe, particularly the Italian honey bee (A. *m. ligustica*) and Carniolan honey bee (A. *m. carnica*), both C-lineage subspecies. Importation of queens has for a long time been used as a means to compensate for the loss of colonies and the Southern European strains are often viewed as a means to improve honey production. It had been assumed that the native UK bee was extinct, but new molecular studies have shown that colonies robustly assigned to *A. m. mellifera* still exist in Northern Europe (13). In the UK the genetic diversity of honey bee populations is poorly understood. The genetic makeup of bee populations not only influences production traits and the ability to survive under less favourable conditions, but also plays a vital role in disease resistance (14). However, the movement of honey bees across the globe has unquestionably contributed to the spread of infectious diseases, which may also be transmitted to wild pollinators (10, 11).

In the UK, the health of honey bees is under threat from a range of native and non-native bacterial, fungal and viral pathogens. While known ‘notifiable diseases’ can be risk assessed and regulated by law, emergent diseases such as *Nosema ceranae* (15) may be spread globally before they have been properly identified and risk assessed. Nosemosis is one of the most prevalent honey bee diseases and is caused by two species of microsporidia, *Nosema apis* and *Nosema ceranae*, that parasitize the ventriculum (midgut). Although infected bees often show no clear symptoms, heavy infections can result in a broad range of detrimental effects (16–21). *N. ceranae*, a native parasite of the Asiatic honey bee (*Apis cerana*), has been detected in *Apis mellifera* samples from Uruguay predating 1990 but is now present in *Apis mellifera* worldwide (15). Notifiable diseases, American foulbrood (AFB) and European foulbrood (EFB), are caused by the non-native bacteria *Paenibacillus larvae* and native *Melissococcus plutonius*, respectively (22, 23). Acarine disease is caused by a mite found throughout Britain which infests the trachea of honey bees (24). Protozoans such as gregarines and the emergent trypanosomatid *Lotmaria passim*, also infect honey bees. The most devastating of all pathogenic species in recent years is the hemophagous mite *Varroa destructor*, which shifted hosts from *A. cerana* to *A. mellifera* sometime in the first half of the 20th century (25). *Varroa* mites feed on the haemolymph of both larval and adult stages of the honey bee. More importantly, *V. destructor* transmits several bee viruses, generating epidemics that kill colonies within two to three years unless the *Varroa* population is kept under control. Among the most important and lethal viruses in this regard are deformed wing virus (DWV) (26), acute bee paralysis virus complex (ABPV), Kashmir bee virus (KBV), and Israeli acute paralysis virus (IAPV) (27). Sacbrood virus (SBV) can also be transmitted but without major epidemic consequences and is primarily indirectly affected by *Varroa* (25, 28, 29).

The core commensal microbiome can mediate disease susceptibility and the internal ecology of the host can greatly affect disease outcome (30). In addition to immunological health and essential nutrient provision, microbial metabolism affects the growth, behaviour and hormonal signalling of honey bees (31). Unlike most host species, the core microbiota of the honey bee has relatively little diversity (32–38). *Snodgrasella alvi* (Betaproteobacteria), *Gilliamella apicola* (Gammaproteobacteria), two *Lactobacillus* taxa (Firm-4 and Firm-5) (34, 35), and *Bifidobacterium asteroides* are common and abundant (39, 40). There are at least four less common species: *Frischella perrara* (41), *Bartonella apis* (42), *Parasaccharibacter apium* (37) and Gluconobacter-related species group Alpha2.1 (35). Metagenomic analyses have revealed high between-isolate genetic diversity in honey bee microbiotal taxa, suggesting they comprise clusters of related taxa (43). These bacteria maintain gut physiochemical conditions and aid their host in the digestion and metabolism of nutrients, neutralisation of toxins, and resistance to parasites (44–46). *Gilliamella* species digest pectin from pollen, and the *Lactobacillus* species inhibit the growth of foulbrood bacteria (47). However, *F. perrara* may cause a widespread scab phenotype in the gut (48). A negative correlation was found between the presence of *Snodgrasella alvi* and pathogenic *Crithidia* in bees (49), but pre-treatment of honey bees with S. *alvi* prior to challenge with *Lotmaria passim* (an *A. mellifera* pathogen closely related to *Crithidia*) resulted in greater levels of *L. passim* compared to bees which were not pre-treated (50). Thus, commensal microbiome species can have beneficial, mutual or parasitic relationships with their hosts, and in particular, different combinations of species – different microbiota communities – may be associated with variations in honey bee health.

With recent significant reductions in the cost of high throughput sequencing, metagenomics could be a useful tool for analysing genetic lineage, gut health and pathogen load as part of routine testing and/or monitoring imports for novel pathogens. Here, to establish baseline figures and test the suitability of this approach, we applied a novel network analysis framework together with deep sequencing of the honey bee metagenome, examining the genomes of honey bees and their symbiotic and pathogenic cobionts in UK apiaries.

## Results

### Metagenome sequencing of honey bees and their cobionts

We performed full metagenomic sequencing of 19 samples of UK honey bees (**Supplementary Table 1**). Samples were obtained from hives located across Scotland and England (**Fig. 1a**), each sample comprising of 16 workers collected from a single colony. Duplicates of samples 1–4 were analysed at a lower sequencing coverage to assess cobiont and genomic variant discovery. While the sample size was limited, the colonies sequenced were selected as representative of the phenotypic diversity of honey bees currently managed by UK apiarists. Notably, representatives of the Buckfast bee and the Colonsay “native” black bee lines were included in the sampling. The entire thorax and abdomen was processed for genome sequencing, thus including gut microorganisms, organisms attached to the outside of the bees, and haemolymph/tissue parasites. Between 4.5 and 12.5 million 125 base paired-end reads were generated per sample on the Illumina HiSeq 2500, equivalent to between 17- and 50-fold coverage of the honey bee genome (Amel 4.5).

**Figure 1:**
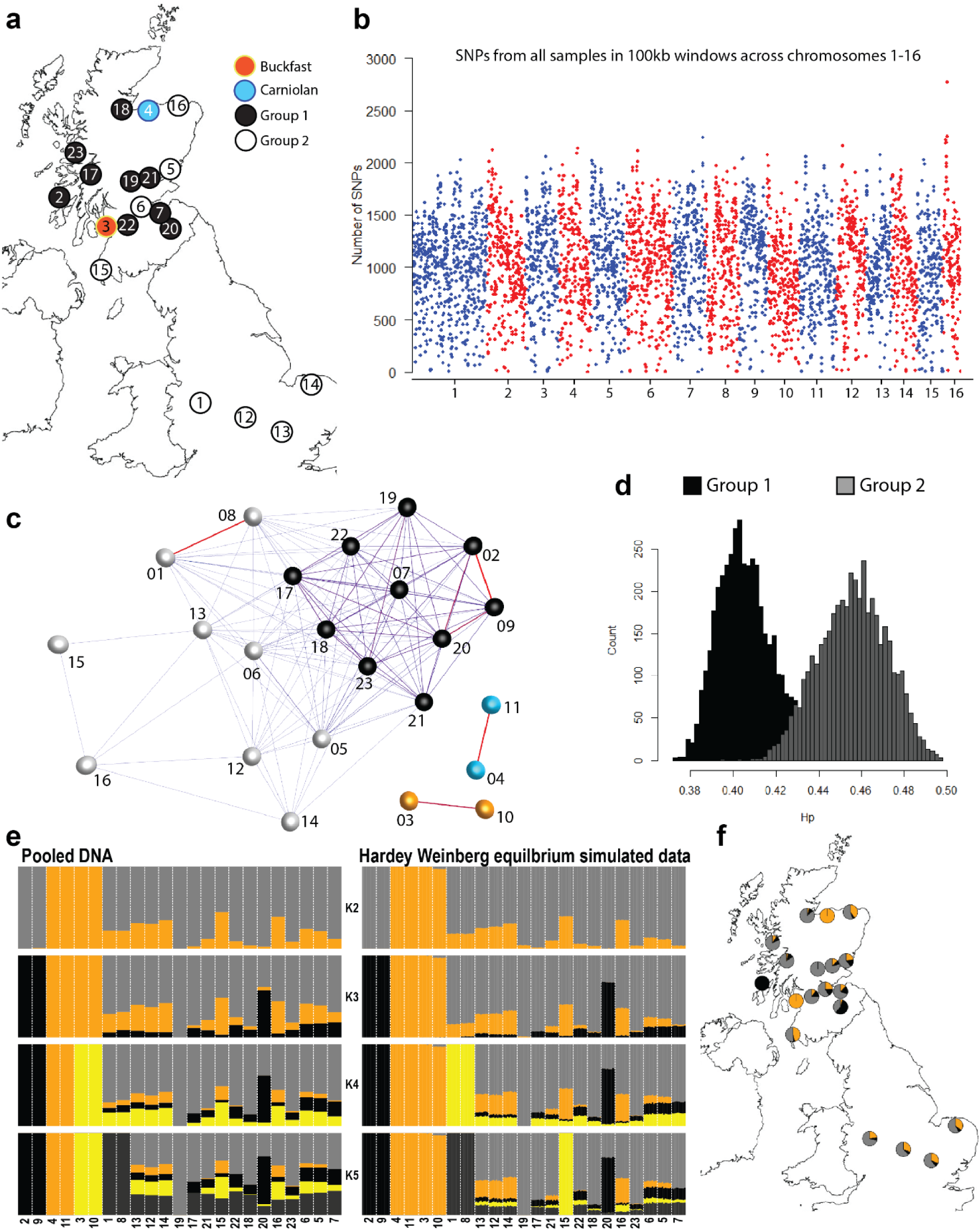
*Apis mellifera* diversity. **(A)** A map of the UK with the location of colonies sampled. **(B)** The number of SNVs from all samples presented across *A. mellifera* chromosomes 1 to 16 in 100 kb consecutive windows. **(C)** A network based on the identity by state (IBS) similarity score of sample variants identifying Groups 1 and 2 in the major cluster. This includes sequencing duplicates (01–04). **(D)** The heterozygosity level across consecutive window of size 100 kb comparing groups 1 and 2 identified from the network graph. **(E)** ADMIXTURE analyses of pooled DNA (left) and genotypes simulated assuming Hardy Weinberg equilibrium (right). **(F) M**ap of sampling locations indicating ADMIXTURE results at K = 3.

### Genomic diversity of sampled honey bees

DNA sequence data were mapped onto the honey bee reference genome (version Amel 4.5 (51)) and variants identified. Overall 3,940,467 sites were called as polymorphic, ranging from 962,775 to 2,586,224 single nucleotide variants (SNVs) per sample (**Fig. 1b**). A correlation graph derived from a matrix of identity-by-state (IBS) at each variant position for all samples was used to define related groups of samples (**Fig. 1c**). Group 1, which includes the native black bee sample from Colonsay (samples 2 and 9), was less heterozygous than Group 2 (**Fig. 1d**). ADMIXTURE (62) analyses were used to explore population subdivision in the data following removal of SNVs in linkage disequilibrium. ADMIXTURE cross-validation (CV) error values increased as the number of populations (K) assumed to be contributing to the variation were increased (K=1, CV= 0.562; K=2, CV= 0.601; K=3, CV= 0.712; K=4, CV= 0.853; K=5, CV= 1.007). At K=2 the Buckfast (samples 3 and 10) and Carniolan (samples 4 and 11) C lineage samples were distinguished from the M lineage *A. m. mellifera* samples, while K=3 further discerns the “native” *A. m. mellifera* sampled from Colonsay (samples 2 and 9), the Buckfast sample at K = 4 and the *A. m. mellifera* breeding project (samples 1 and 8) at K = 5 (**Fig. 1e**).

ADMIXTURE was originally designed to estimate ancestry in unrelated individuals rather than pooled DNA from several individuals, as analysed here. To address this, genotypes were simulated for 10 individuals per pooled DNA sample, using allele sequence depth to estimate allele frequency under an assumption of Hardy-Weinberg equilibrium and analysed using ADMIXTURE. The CV error values decreased as K was increased (K=1, CV= 0.980; K=2, CV= 0.835; K=3, CV= 0.795; K=4, CV= 0.763; K=5, CV= 0.736). At K≤3 the simulated data results were consistent with those from the actual pooled genotypes, while K=4 distinguished samples from the *A. m. mellifera* breeding project (samples 1 and 8), and K=5 assigned a distinct genetic background to bees sampled from Wigtownshire (sample 15) (**Fig. 1e**). k-nearest neighbour (kNN) network analysis of the pooled genotype data (63) also identified 2 clusters, separating C and M lineage samples in the same manner as the ADMIXTURE analyses. Together, these results support a model of two genetic backgrounds in the UK bee populations sampled, most likely representing the C and M lineages, with evidence of a distinct *A. m. mellifera* background in bees originating from Colonsay and other areas of Scotland, and differentiation of Buckfast and Carniolan bees (**Fig. 1f**).

### The microbiome of honey bees

The majority of the data (~90% of reads) from each sample mapped to the honey bee reference genome. Reads that did not map to the honey bee reference were collated and used for a metagenomic assembly. This resulted in over 35,000 contigs greater than 1 kb in length. Contigs were assigned to a taxonomic group by comparison to a series of curated databases in a defined order (**Fig. 2a**) using BlobTools (52). First, contigs were compared to the bee cobiont sequence data in the HoloBee Database (v2016.1) (53), followed by genomes and proteomes of species identified as being bee-associated (54, 55), and finally by comparison of contigs against the NCBI Nucleotide and UniProt Reference Proteome databases. Patterns of coverage, GC% and taxonomic annotation of contigs were explored to identify likely genomic compartments present (**Fig. 2b,c**). We discarded contigs with read coverage lower than 1, as these were likely an artefact of pooling reads, yielding a final set 31,386 metagenome contigs, spanning 140 Mb. Taxon assignments are summarised in **Supplementary Table 2**. Correlation graphs were constructed based on the coverage or contribution to each contig from each sample. Clustering samples in this manner did not recapitulate their clustering by honey bee genome SNVs (**Fig. 2d**). A correlation graph was also constructed where nodes represented individual contigs (**Fig. 3**). A high correlation threshold (*r* = 0.99) was used, which meant that 35% of the contigs were unconnected. The highly structured multi-component graph was subdivided using the MCL algorithm (56) into clusters of contigs whose abundance across the samples was very similar. Many of these clusters were made up of contigs derived from the same species or in a number of cases from strongly co-occurring species.

**Figure 2:**
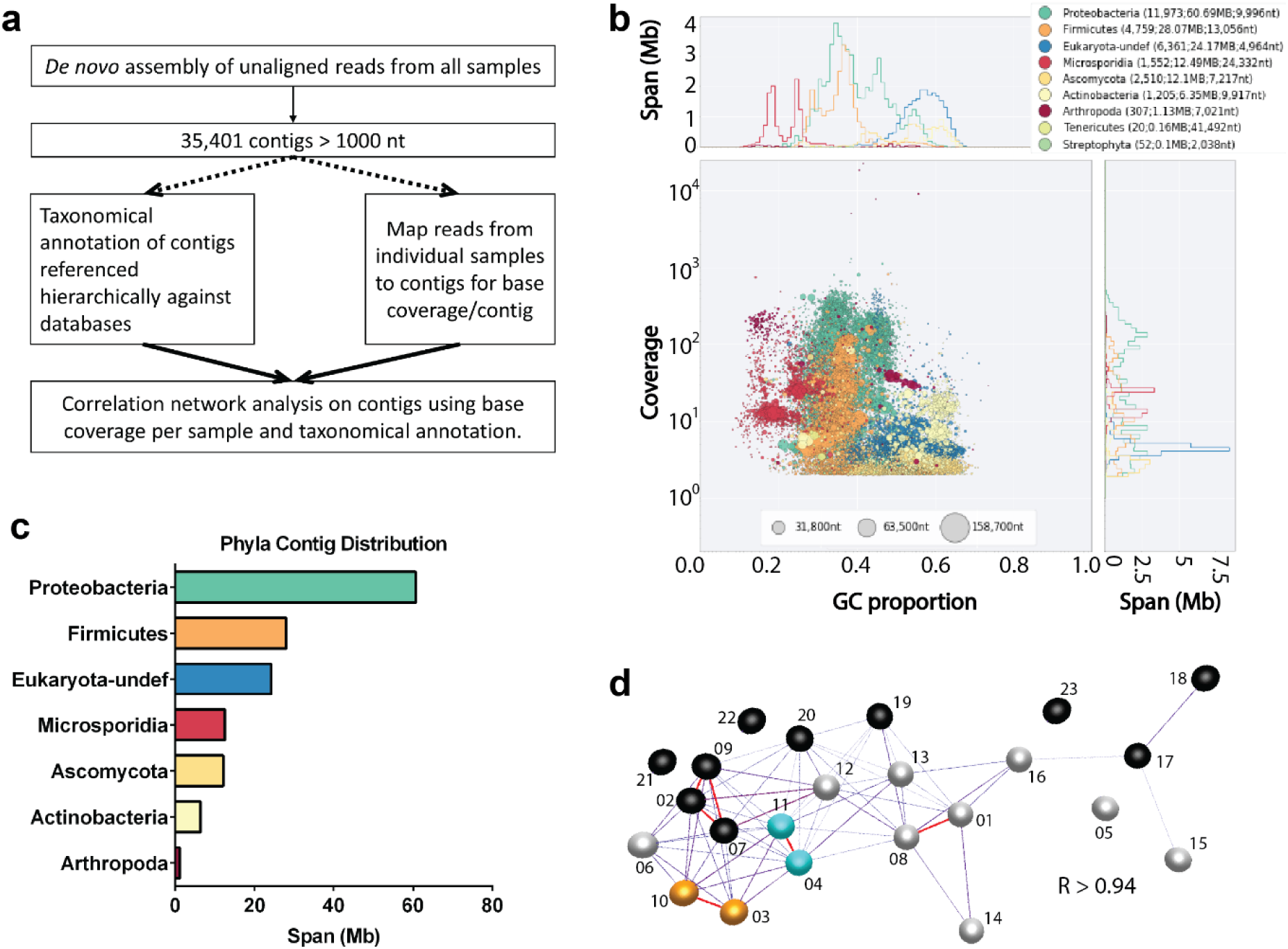
Metagenomics of *Apis mellifera*. **(A)** A flow diagram of the microbiome analysis using reads which did not align to the *Apis mellifera* reference genome. **(B)** A blobplot generated from contigs using unaligned reads from all samples. Contigs are plotted based on their GC content (x-axis) and coverage (y-axis), scaled by span, and coloured by their phylum assignation. **(C)** The span of *de novo* assembled contigs which were assigned to given phyla is displayed for the 12 most abundant phyla across all samples. (D) A network based on the coverage/contig from each sample representing microbiome composition/unaligned reads.

**Figure 3:**
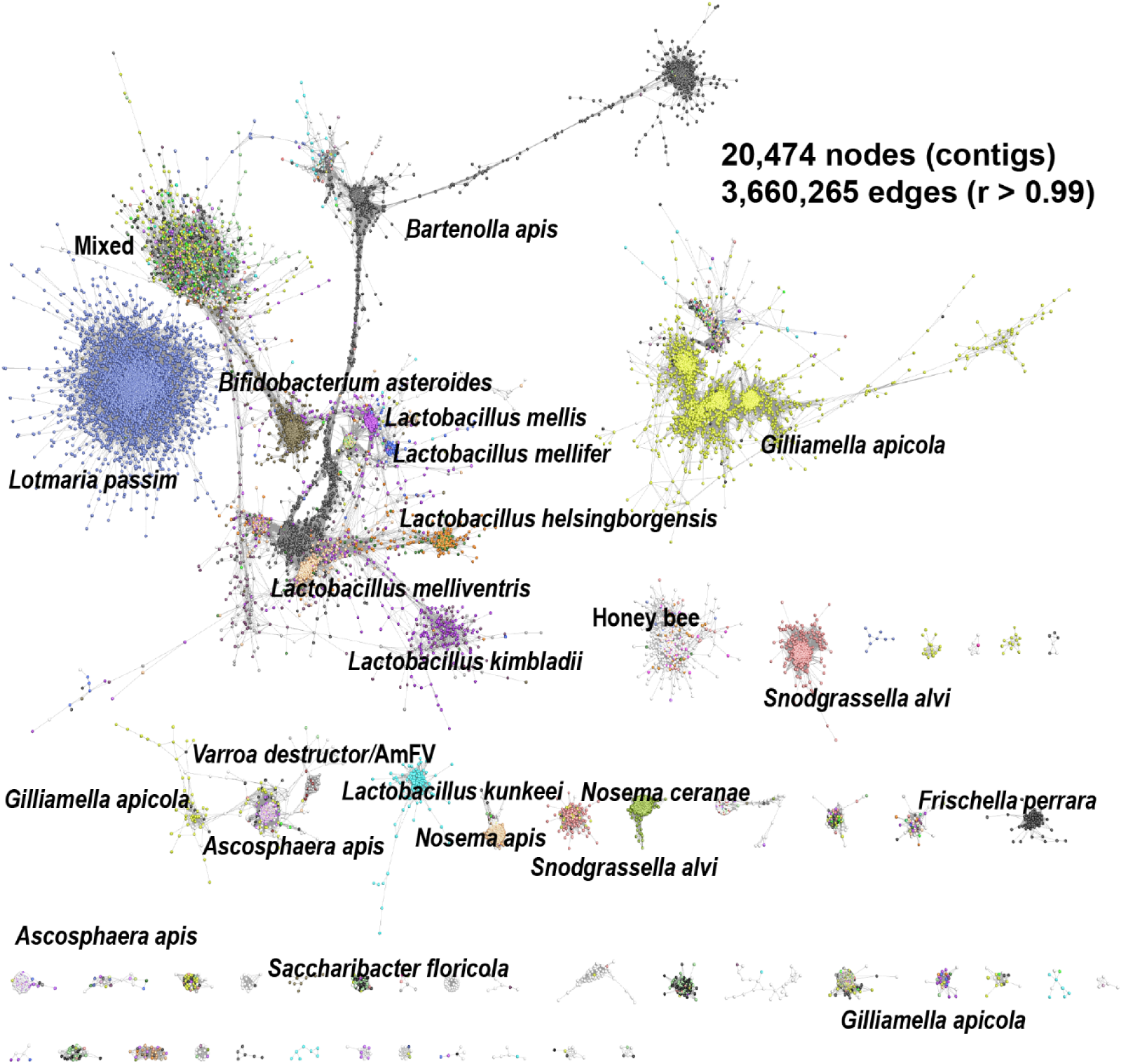
Correlation network analysis of microbiome contigs. Each node represents an individual contig and edges are defined based on the abundance profile of the number of reads mapping to the contig across individual samples. Contigs (nodes) are connected if the Pearson correlation between two contigs abundance profile was *r* > 0.99. Each contig is coloured according to species ID, white nodes represent contigs for which no significant sequence match was found.

Rarefaction analysis of ribosomal RNA sequences present in the assembled data was used to estimate the species richness discovered as a function of sequencing depth (**Supplementary Fig. 1**). While there was variation between samples in terms of species richness at all sequencing depths, even the lowest coverage achieved (17x reference genome coverage) was likely to be sufficient to capture most *A. mellifera* cobionts present.

We examined clusters on the graph further. One (**Fig. 4a**) contained 1.33 Mb of sequence, most of which had no match in public databases, but contained some contigs that had significant similarity to sequences from other *Apis* species (**Fig. 4b**). The number of reads mapping to these contigs was proportional to the depth of honey bee genome sequencing (**Fig. 4c**) and we infer that they likely represent reads from true *A. mellifera* genome fragments not present in the honey bee reference genome (**Fig. 4d**). Others in this cluster, spanning 0.01 Mb, matched sequences from *Ascophaera apis* (chalkbrood), an endemic fungal associate of honey bees (57).

**Figure 4:**
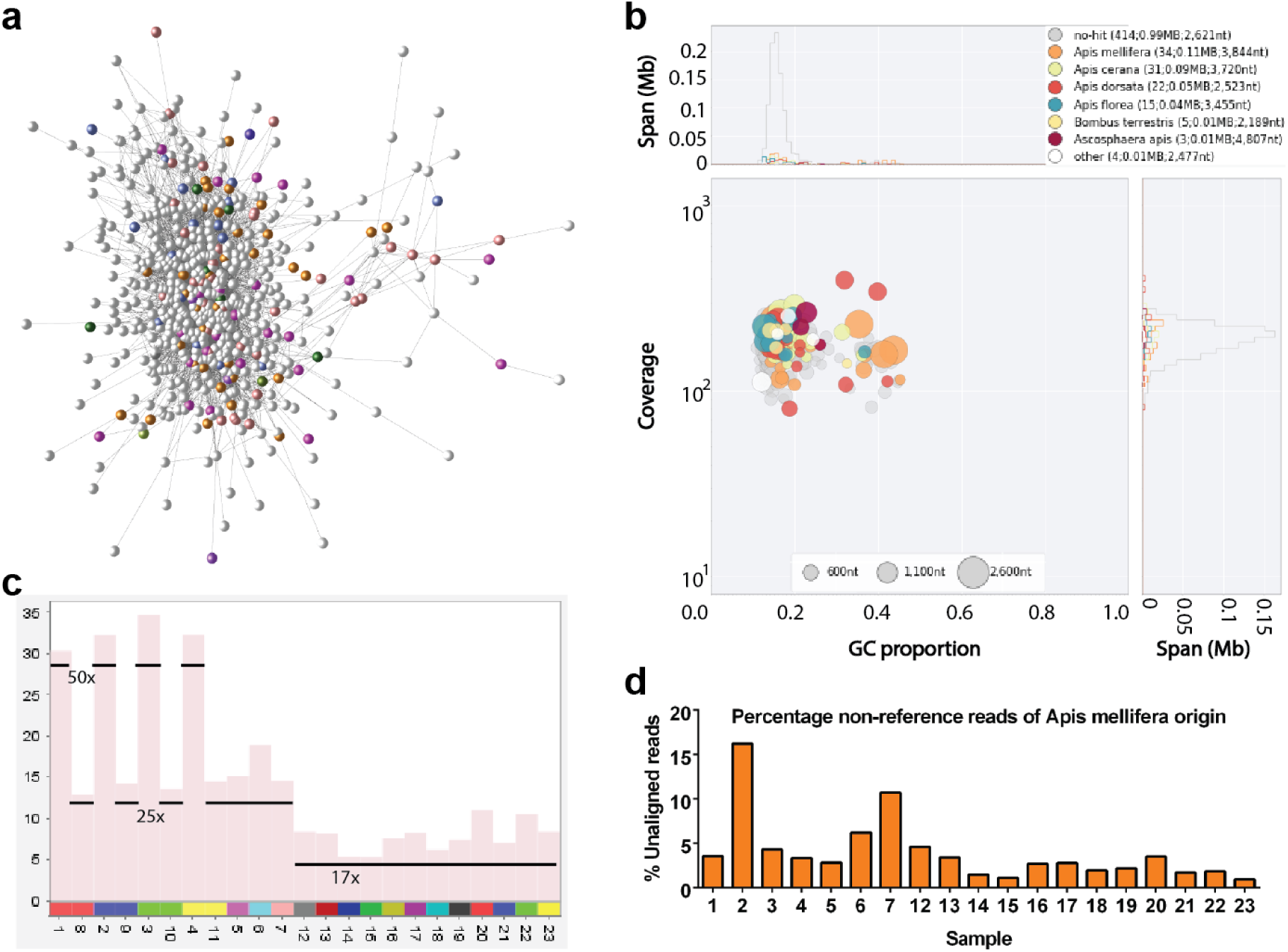
Putative *Apis mellifera* contigs. **(A)** Network component of contigs which did not match the reference bee genome and are unassigned (white) or match a non-reference species of bee (coloured). **(B)** Blobplot of these contigs (as in Figure 2). **(C)** Mean base coverage per contig (y-axis) for each sample (x-axis) for the contigs in A. The sequencing depth (reference genome coverage) per sample is shown, showing that the number of reads mapping to these contigs is in direct proportional to the depth of sequencing. **(D)** A graph displaying the percentage of unaligned reads putatively identified as *Apis mellifera* from each sample.

Most of the other groups of contigs could be assigned to cobiont organisms. The contribution of non-*A mellifera* reads varied between samples, a pattern that may be partly explained by the presence in some samples of eukaryotic pathogens such as *Nosema* microsporidians and the trypanosomatid *L. passim*, which have larger genomes. The most abundant non-pathogenic bacterial cobionts identified were *Gilliamella apicola, Bartonella apis, Frischella perrara, Snodgrassella alvi*, “Firm-4” firmicutes (58) (*Lactobacillusmellis* and *Lactobacillusmellifer*), “Firm-5” firmicutes (58) (*Lactobacillus melliventris, Lactobacillus kimbladii, Lactobacillus kullabergensis, Lactobacillus sp. wkB8, Lactobacillus helsingborgensis* and *Lactobacillus sp. wkB10*)*, Lactobacillus kunkeei* and *Bifidobacterium asteroides* (**Supplementary Table 2**). Each species varied in its abundance across the samples. In some nominal species, contig read coverage clustering suggested the presence of multiple distinct genotypes of cobionts. Contigs ascribed to *Bartonella apis* together had a total span of 11.7 Mb, almost five times longer than the reference *B. apis* genome, and formed a connected module (**Fig. 5a**). The three largest *B. apis* clusters had distinct distribution across the samples, which we suggest reflects the presence of distinct genotypes of *B. apis* with varying abundance across the samples. Similarly, contigs ascribed to *Gilliamella apicola*, the most abundant species identified in the bee microbiome, were distributed across three closely related clusters (**Fig. 5b**). Groups containing contigs from several closely related but distinct *Lactobacillus* species were identified: Firm-4 lactobacilli (clusters 25 and 40) or Firm-5 lactobacilli (clusters 16, 20, 21 and 24) (**Fig. 5c**). These *Lactobacillus* groups may represent distinct cobiont communities. The exception was cluster 21, which contained contigs assigned to a mix of Firm-5 species: this may represent a core genome component conserved between species. Cluster 29 comprised contigs assigned to *Lactobacillus kunkeei* that formed an unconnected graph component. *L. kunkeei* is thought to be an environmental rather than a gut microbiome organism. Some connected components were more complex. Cluster 32 contained contigs assigned to several prevalent honey bee cobionts, including *G. apicola, F. perrara, B. asteroides, S. alvi, B. apis, S. floricola* and *P. apium*. The co-clustering of genomic segments from multiple species is likely to reflect a strongly interacting community of organisms where the relative abundance of each is regulated homeostatically (43, 55, 59).

**Figure 5:**
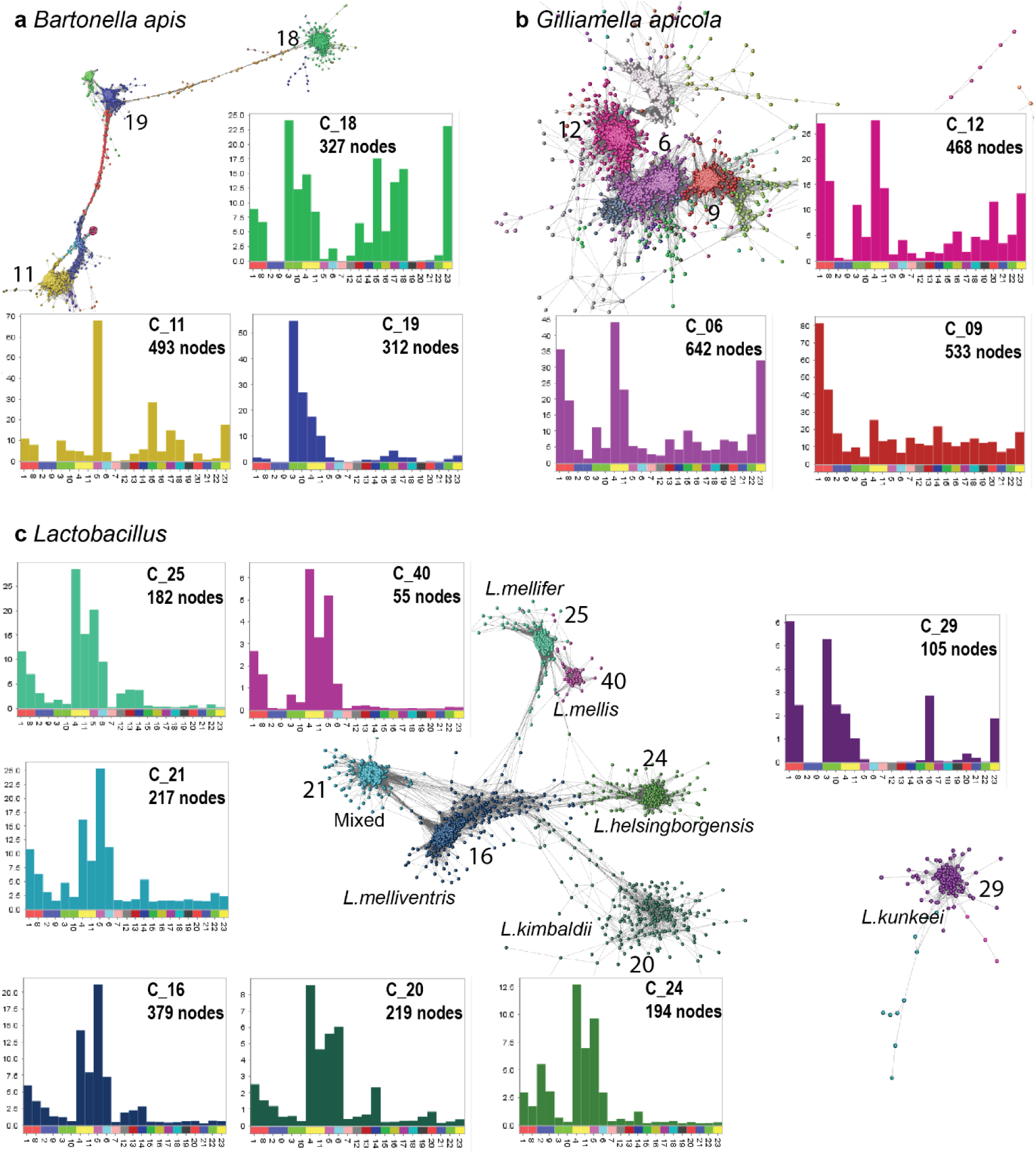
Communities of honey bee cobionts. Sub-networks of contig clusters from Figure 3 coloured by cluster. Histograms show the mean base coverage per contig (y-axis) for each sample (x-axis). **(A)** *Bartonella apis*, **(B)** *Gilliamella apicola* and **(C)** several *Lactobacillus* species. Blobplots describing the taxonomy and cumulative span for each of these panels are presented in **Supplementary Figure 1A-C**.

Some clusters had very restricted presence in the sample set. For example, cluster 3 was largely restricted to sample 4 (**Supplementary Fig. 2e**). These are likely to derive either from rare members of the honey bee cobiont community or opportunistic infections. Several clusters had little to no annotation (**Supplementary Fig. 2f**). The coverage of these contigs was also usually derived from individual samples. They may represent novel species, or divergent or novel genomic regions of known species.

### Honey bee pathogens

Known honey bee pathogens were detected in many samples. One of the largest components of clustered contigs was assigned to the trypanosomatid parasite *Lotmaria passim*, with a combined span of 16.3 Mb (**Fig. 6a**). While sequences were detected from notifiable pathogens *Melisococcus plutonious* and *Paenibacillus larvae* (European and American foulbrood), no distinct cluster was identified and the <1Mb total combined span of matched sequences was relatively minor (**Supplementary Table 2**).

**Figure 6:**
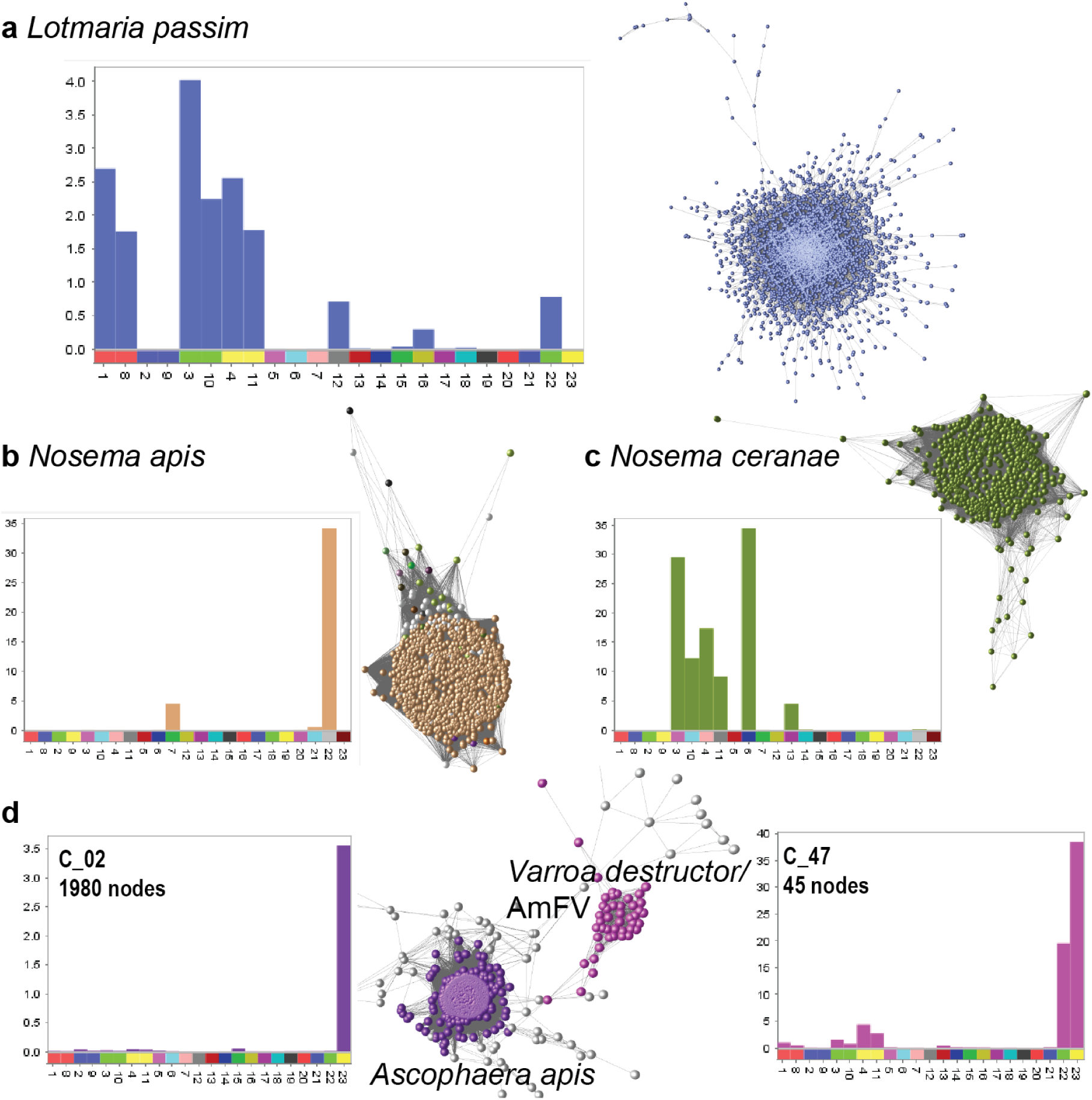
Disease associated components. Clusters associated with honey bee cobionts including mean base coverage per contig (y-axis) for each sample (x-axis). **(A)** *Lotmaria passim*, **(B)** *Nosema apis*, **(C)** *Nosema ceranae* and **(D)** a community of species including *Ascophaera apis* (associated with chalkbrood), *Varroa destructor* and *Apis mellifera filamentous virus* (AmFV). Blobplots describing the taxonomy and cumulative span for each panel are presented in Supplementary Figure 1D-J.

Both *Nosema* species *N. apis* (**Fig. 6b**) and *N. ceranae* (**Fig. 6c**) were identified. *N. ceranae* was more prevalent (4/19 colonies vs. 2/19 colonies). Contigs matching the pathogen causing “chalk brood” (*Ascophaera apis*) were found in cluster 2 and were derived almost exclusively from sample 23 (**Fig. 6d**). In close proximity in the network graph was cluster 47, containing contigs assigned to the parasitic mite *V. destructor* and contigs assigned to *Apis mellifera filamentous virus* (AmFV), found in 6/19 colonies (**Fig. 6c**). The largest source of reads mapping to these contigs was sample 23, which also had a high prevalence of chalkbrood. Blobplots describing the taxonomy and cumulative span for each panel in **Fig. 6** are available in **Supplementary Fig. 2d-j**.

To validate the metagenomic hits, we employed PCR to screen our samples for *B. apis, Nosema ceranae* and *L. passim*. All samples in which we identified sequences deriving from organisms were positive by their respective PCR. However, we also identified the presence of species in additional samples not scored as positive by sequencing, suggesting that the PCR assays are more sensitive than bulk sequencing (**Supplementary Fig. 3a-c**). We also identified a small cluster containing only one contig matching to a recorded genome sequence, *Apicystis bombi*, a gregarine known to parasatise honey bees (60). To identify the exact species present, we sequenced the PCR results of custom primers against the largest contig in this cluster, in conjunction with primers encompassing the 18 S and ITS2 rDNA regions, as used by Dias *et al*. for the characterisation of novel gregarine species (61) (**Supplementary Fig. 3d**). The contig sequence matched with various gregarine species, while the ribosomal DNA sequence confirmed the species present to be *Apicystis bombi* (**Supplementary Fig. 3e**).

## Discussion

A healthy population of honey bees is crucial for the security of the ecosystem service of pollination. With the continued and sometimes unregulated global transport of *A. mellifera*, the introduction of invasive pests and parasites is a continuing threat, as is the genetic dilution or extinction of locally adapted subspecies. Here we used metagenomic analyses of nineteen honey bee colonies from around the UK to compare host genetics, examine the complexity and connectedness of the bee microbiome, and quantify disease burden.

Using the reference honey bee genome and sequence data from sixteen worker bees from each colony, we defined over five million SNVs with a relatively even distribution across all sixteen chromosomes (**Fig. 1**). We also identified likely honey bee-derived sequences not represented in the reference genome, likely because the reference is incomplete or because of variation in genomic content between honey bee populations. The island of Colonsay in Scotland is a reserve for the northern European bee, *A. m. melifera*. Given the level of bee imports into Scotland, it was therefore reassuring – and perhaps surprising – to observe that the genotypes of other colonies from around Scotland were close to that of the Colonsay sample, although distinct from samples from *A. m. mellifera* breeding programmes in England. The low heterozygosity of Scottish *A. m. mellifera* and continued survival in face of imports may reflect natural selection for *A. m. mellifera* genotypes in the colder climates and shorter foraging season of northern Europe.

The whole organism-derived sequence data was also used to explore the composition of the communities of organisms living in or on honey bees. Non-*A. Mellifera-mapping* reads were assembled to generate 160 Mb of genomic sequence from honey bee cobionts. These cobionts were biologically identified using read depth coverage, patterns of coverage across samples and best taxonomic assignment based on comparisons to known organisms. A correlation network based on per-sample read coverages of these contigs (**Fig. 2d**) did not fully match the relatedness of the source bees (**Fig. 1c**), suggesting that both environmental and host genetic components drive microbiome composition. Our limited sampling (only nineteen colonies) is not sufficient to unpick these interdependent drivers, but we note that samples from the Scottish coast, the central belt of Scotland and from England were grouped separately. These data are congruent with previous analyses of the roles of climate and forage in determining microbiome structure of honey bees (62, 63).

In many animals, the gut microbiota form quasi-stable communities, with individual hosts harbouring somewhat predictable communities of different bacterial taxa. These different microbiome types have been associated with different gross physiological performance. In addition, changes in microbiota composition (dysbiosis) have been associated with the promotion of disease states in humans and other mammals (64, 65). Dysbiosis in honey bees may be an important correlate of bee and colony health (48, 66–68).

In the honey bee gut, bacterial numbers are highest in the rectum, followed by the ileum, mid-gut and crop (66). Lactobacilli are mainly found close to the rectum and, together with bifidobacteria, greatly outnumber other species (66). We identified several contig clusters that likely represented single *Lactobacillus* species as well as a mixed-origin cluster (**Fig. 4**). Most of these were interlinked, revealing patterns of co-occurence of individual taxa. In contrast, *L. kunkeei*, an environmental cobiont reportedly indicative of poor health (66), formed a distinct, unlinked cluster. Samples 2 and 9 were technical replicates, and both had reduced diversity, containing only *G. apicola* and *Lactobacillus* species. The reason for this is unclear, but there was no evidence of pathogenic disruption of the sampled bees.

*Nosema* infection has been linked to immune suppression and oxidative stress of bee hosts (69). Similarly *L. kunkeei* and *P. apium*, which are adapted to fluctuating oxygen levels predicted for the gut (70), have been associated with disease states in social bees, and negatively correlated with the amount of core commensal bacteria present (66). The microbiome from sample 23 was had a preponderance of reads mapping to the *L. kunkeei* cluster (**Supplementary Fig. 2c**), evidence of *P. apium* presence, much reduced representation of other *Lactobacillus* species, and the highest read coverage of contigs associated with the pathogens *V. destructor*, AmFV and *A. apis*. Sample 23 may be an example of pathogen-induced dysbiosis, or of invasion by pathogens of a resident microbiome disturbed by other drivers. There was a high level of co-occurrence of different pathogens across samples, implying that colonies infected with one pathogen may be more susceptible to others. A meta-stable community may exist in the case of *Varria destructor* and AmFV (**Fig. 6c**). However, we note a recent study reported identifying 0.5 Mb of sequence from *Varroa* reference genome to be of AmFV origin (71). It is therefore possible that several of the contigs in our study matched with *Varroa destructor* are in fact of AmFV origin.

Several distinct contig clusters were assigned to *G. apicola* and *B. apis* suggesting the existence of genetically distinct subtypes of these highly prevalent bacteria. (**Fig. 5a,b**). *G. apicola* has a high diversity of accessory genes, associated with adaptation to different *A. mellifera* ecological niches (72, 73). Increased relative abundance of *G. apicola* has been associated with dysbiosis and host deficiencies (66). Similarly, extreme displacement of S. *alvi* by *F. perrara* and *G. apicola* (and to a lesser extent by the opportunists *P. apium* and *L. kunkeei*) has been strongly associated with reduced bacterial biofilm function and host tissue disruption by scab-inducing *F. perrara* (48, 68), leading to poor host development and early mortality. Blooms of *B. apis* have also been associated with poor health. This species exploits stressed, young, and old bees, showing sporadic abundance in whole guts of newly emerged workers (58) and occurring uniformly across putatively dysbiotic foragers (56). In support of this theory, samples from our study with the highest coverage of *G. apicola* and *B. apis* contigs also contained reads from pathogens such as *L. passim* or *Nosema* species. Significant positive correlation has been reported between infection levels of these parasites (74).

Our novel use of correlation networks (**Fig. 3**) to organise contigs based on their relative abundance across samples partitioned 65% of them into clusters of sequences corresponding to individual species and distinct micro-communities. Some sample-specific clusters, such as clusters 3 and 32, contained several core microbiome taxa. This may be a reflection of substrate specialisation based on host foraging (75). However, several sample specific clusters contained contigs that had no informative taxonomic annotation, potentially revealing uncharacterised species. We identified a cluster of unclassified contigs derived from a gregarine, with closest match to *Apicystis bombi*. The accuracy of our metagenomic analyses was confirmed by PCR and ribosomal DNA primers verified the species as *Apicystis bombi*. This is further evidence that managed honey bees can act as a reservoir for wild pollinator pathogens (60); through increased understanding of honey bee molecular ecology and preventing disease transmission, we can indirectly improve wild pollinator health (76). To our knowledge *Lotmaria passim* had not been previously identified in the UK. Its presence was confirmed for the first time in our study using the primers designed by Stevanovic *et al*. (77), further validating our sequencing inference.

A whole-organism metagenomics approach has allowed us to describe the complexity of host-microbiome biology of UK honey bees. Using pooled samples we have demonstrated the power of this approach in dual characterisation of the genotypic diversity of the honey bee and the genomic diversity of its cobionts. Correlation networks are powerful analytic tools that allowed us to cluster the sequence data to reveal interacting networks of bacterial and eukaryotic microbiota, in addition to classifying novel genomic sequences. As with the human and other animal microbiome projects, the precision of these analyses improves with additional data, permitting definition (and ultimately whole genome assembly) of novel genotypes of cobionts. To this end, the raw data from this project can be accessed through the Bee Microbiome Database, established and managed by the Bee Microbiome Consortium, a non-profit organization of bee scientists for collecting, curating and analysing bee microbiome data (55). Complementation of cheap short read data with low-coverage long-read data from isolated gut contents enhances the contiguity of assemblies and the functional inferences that can be derived them. This study highlights the potential to use this approach in routine screening, breeding programmes and horizon scanning for emerging pathogens.

## Methods

### Samples

Nineteen samples of honey bee (each comprising sixteen workers collected from a single colony) were obtained from beekeepers in Scotland and England, with the help of Science and Advice for Scottish Agriculture (SASA) and Fera Science Ltd. Wings, legs and heads were removed before homogenizing the remainder of the bees (thorax and abdomen) in 2% CTAB buffer (100 mM Tris-HCl pH 8.0, 1.4 M NaCl, 20 mM EDTA pH 8.0, 2% hexadecyltrimethylammonium bromide, 0.2% 2-mercaptoethanol). Samples were incubated at 60ΰC with proteinase K (54 ng/μl) for 16 h before incubating with RNaseA (2.7 ng/μl) at 37ΰC for 1 h. After two chloroform:isoamyl alcohol (24:1) extractions, samples were ethanol precipitated, washed three times in 70% ethanol and resuspended in 0.1 TE. All genomic DNA samples were analysed for quantity (Qubit dsDNA HS Assay Kit, Thermo Fisher Scientific, Waltham, MA, USA), purity (Nanodrop, Thermo Fisher Scientific, Waltham, MA, USA) and quality (TapeStation, Agilent Technologies, Santa Clara, CA, USA).

### Sequencing

All sequencing was performed by Edinburgh Genomics. DNAs were prepared for whole genome sequencing using the TruSeq DNA PCR-free gel free library kit (Illumina, Cambridge, UK) and, for eight samples, using the TruSeq DNA Nano gel free library kits (Illumina). For comparison, both types of libraries were prepared for four samples. 125 base paired-end sequencing was performed on an Illumina HiSeq 2500. Four samples were sequenced at 50X coverage, eight at 25X (including repeat sequencing of the four 50X samples) and twelve at 17X coverage. Data were screened for quality using FastQC v0.11.2 (Available online at: http://www.bioinformatics.babraham.ac.uk/projects/fastqc), and trimmed of low quality regions and adapters using Trimmomatic v0.35 (78) with parameters ‘TRAILING:20 SLIDINGWINDOW:4:20 MINLEN:100.’ These parameters remove bases from the end of a read if they are below a Phred score of 20, clip the read if the average Phred score within a 4 base sliding window advanced from the 5’ end falls below 20, and specify a minimum read length of 100 bases (the parameters used for all informatics analyses are also detailed in **Supplementary Table 3**).

### Variant calling on honey bee

Reads were aligned to the reference *A. mellifera* genome, Amel_4.5 (INSDC assembly GCA_000002195.1) using BWA-MEM v0.7.8 (79) with parameters -R and -M. Output files were merged and duplicates marked using Picard Tools v2.1.1 to create one BAM file per sample. This was filtered using SAMtools view v1.3 (80) to retain only the highest confidence alignments using the parameters -q 20 (to remove alignments with a Phred score <20) and -F 12 (to remove all reads that are not mapped and whose mate is not mapped).

Variants were called using GATK v3.5 in accordance with GATK best practice recommendations (81, 82). Local realignments were performed and base quality scores recalibrated using bee SNVs from dbSNP (83) build ID 140 (ftp://ftp.ncbi.nlm.nih.gov/snp/organisms/bee_7460/VCF/, downloaded 1^st^ January 2016). GATK HaplotypeCaller was used with parameters emitRefConfidence, - GVCF variant index type – LINEAR, variant index parameter -128000, stand emit conf – 30, stand call conf - 30. The resulting VCFs, one per sample, were merged to create a single gVCF file using GATK GenotypeGVCFs to allow variants to be called on all samples simultaneously. Variant quality score recalibration was performed on this file using GATK VariantRecalibrator with parameters badLodCutoff – 3, -an QD, -an MQ, -an MQRankSum, -an ReadPosRankSum, -an FS, -an DP (specifying the above dbSNP data as both the truth set [prior=15.0] and training set [prior=12.0]). To identify any effect these variants may have upon protein-coding genes in the reference annotation, we used SNPeff v4.2 (84). A total of 5,302,201 variants were identified across the 19 samples.

An Identity By State (IBS) analysis was performed using the R/Bioconductor package, SNPRelate (85). Briefly, the minor allele frequency and missing rate for each SNV was calculated over all the samples. The values of the resultant IBS matrix ranged from zero to one. Using this matrix, we constructed a network correlation graph for all of the samples, using the network analysis tool Graphia Professional (Kajeka Ltd., Edinburgh, UK), where each node represented a sample, and edges between nodes represented a correlation above the defined threshold between those samples.

### *De novo* assembly and analysis of non-honey bee data

*De novo* assembly was performed on all of the reads which did not map to the *Apis mellifera* reference genome using SPAdes v3.8.1 (86). The resulting contigs were filtered by length (> 1kb) and coverage (> 2). BWA-MEM (79) was used to identify and remove reads mapping to these contigs and *de novo* assembly was performed on the remaining reads. This process was repeated for a total of five iterations. Input reads from each sample were mapped to each contig using BWA-MEM and base coverage/contig was calculated. Contigs with a cumulative base coverage from all samples less than half the SPAdes overall coverage were discarded. Using BLAST (87), contigs were compared to a set of custom databases: 1. HB_Bar_v2016.1 (53); 1. HB_Mop_v2016.1 (53); 3. nucleotide sequences of core microbiome species identified from literature (43, 44, 55, 73); 4. protein sequences of these species (43, 44, 55, 73); 5. NCBI nt (88); 6. UniProt Reference Proteomes (89) using BLAST (87) and Diamond (90). Files of all six sequence similarity searches were provided as input to BlobTools in the listed order under the tax-rule ‘bestsumorder’, *i.e*. a contig is assigned the NCBI taxid of the taxon providing the best scoring hits within a given file, as long as it has not been allocated a NCBI taxid in a previous file.. BlobTools was used to visualise the coverage, GC% and best BLAST similarity match of the assembly, and to build a table of base coverage of contigs in each sample together with their taxonomic annotation. A network graph was constructed using *r* value of 0.99 comparing samples to each other based on correlations between their overall microbiome content as well as contig coverage across the dataset. This follows the approach used to compare gene expression values in transcriptomics data (91).

### Population genetics analyses

SNVs were filtered using Plink v1.9; (92) to remove those not mapped to the autosomes, those having low genotyping call rate (<0.9), those with low minor allele frequency (<0.1), and those with pairwise linkage disequilibrium r^2^ > 0.1 (for SNVs in 50 kb windows with a 10 kb step). The resulting 58,354 SNVs were submitted to unsupervised analyses in ADMIXTURE (93) for 1 ≤ K ≤ 5 genetic backgrounds. To explore consequence of analysing genotypes from pooled DNA, individual genotypes simulated for 10 individuals per sampling location for each SNV were subjected to ADMIXTURE analysis. Briefly, for each SNV the allele frequency observed in a pooled sample was calculated from the read counts for each allele, and used to simulate ten genotypes assuming Hardy-Weinberg equilibrium. The efficacy of this process was tested using data from Harpur *et al*. (94), details of which are provided in the supplementary information (**Supplementary Table 3**). A distance matrix from the pooled DNA genotypes used in ADMIXTURE analyses was generated with Plink and analysed using the R package netview (95) (https://github.com/esteinig/netview), which analyses genetic structure using mutual k-nearest neighbour (kNN) graphs. Graphs were created assuming 2 ≤ k ≤ 20 nearest neighbours.

### Primer design for identification of cobionts using PCR

Custom primers were designed against the longest contigs we generated matching *Bartonella apis* (Bartonella_Fw 5’-CAGCAGCGCTTATTCCGTTC-3’, Bartonella_Rv 5’-AGTCACGAGCAACAATCGGT-3’) and the Gregarine species (Gregarine_F 5’-GACCACCGTCCTGCTGTTTA-3’, Gregarine_R 5’-GAGGTATCGGGTGCCATGA-3’). Primers were run through NCBI BLAST to confirm specificity (87). *Apicystis bombi* specific primers were used as described in Dias *et al*. (61). Specific primers against *Nosema ceranae* were used as described by Chen *et al*. (96) and *Lotmaria passim* specific primers were used as described by Stevanovic *et al*. (77).

### Rarefaction analysis of microbiome sampling

“Mean species richness” was calculated using the R package ‘vegan’ (97) for each sample at each of the sequencing depths used. Assembled contigs were analysed against the SILVA rDNA (16S and 18S) databases (98) instead of the NCBI nt database to assess species composition. Each contig identified as being from a unique species was counted as one “count” or incidence of discovering that species in the sample.

## References

1. Klein AM, Vaissiere BE, Cane JH, Steffan-Dewenter I, Cunningham SA, Kremen C, et al. Importance of pollinators in changing landscapes for world crops. Proceedings Biological sciences. 2007;274(1608):303–13.

2. Stathers R. The Bee and the Stockmarket: An overview of pollinator decline and its economic corporate significance. Schroders. 2014.

3. Hoehn P, Tscharntke T, Tylianakis JM, Steffan-Dewenter I. Functional group diversity of bee pollinators increases crop yield. P Roy Soc B-Biol Sci. 2008;275(1648):2283–91.

4. Kleijn D, Winfree R, Bartomeus I, Carvalheiro LG, Henry M, Isaacs R, et al. Delivery of crop pollination services is an insufficient argument for wild pollinator conservation. Nat Commun. 2015;6.

5. Potts SG, Imperatriz-Fonseca V, Ngo HT, Biesmeijer JC, Breeze TD, Dicks LV, et al. Summary for policymakers of the thematic assessment on pollinators, pollination and food production. Biota Neotrop. 2016;16(1).

6. Aizen MA, Harder LD. The global stock of domesticated honey bees is growing slower than agricultural demand for pollination. Current biology: CB. 2009;19(11):915–8.

7. Memmott J, Craze PG, Waser NM, Price MV. Global warming and the disruption of plant-pollinator interactions. Ecology letters. 2007;10(8):710–7.

8. Ricketts TH, Regetz J, Steffan-Dewenter I, Cunningham SA, Kremen C, Bogdanski A, et al. Landscape effects on crop pollination services: are there general patterns? Ecology letters. 2008;11(5):499–515.

9. Winfree R, Aguilar R, Vazquez DP, LeBuhn G, Aizen MA. A meta-analysis of bees’ responses to anthropogenic disturbance. Ecology. 2009;90(8):2068–76.

10. Furst MA, McMahon DP, Osborne JL, Paxton RJ, Brown MJF. Disease associations between honeybees and bumblebees as a threat to wild pollinators. Nature. 2014;506(7488):364–+.

11. McMahon DP, Furst MA, Caspar J, Theodorou P, Brown MJF, Paxton RJ. A sting in the spit: widespread cross-infection of multiple RNA viruses across wild and managed bees. J Anim Ecol. 2015;84(3):615–24.

12. Neumann PC, N.L. Honey Bee Colony Losses. Journal of Apicultural Research. 2010;49(1):1–6.

13. Bouga M AC, Bienkowska M, Buchler R, Carreck NL, Cauia E, Chlebo R, Dahle B, Dall’Oli, R, De la Rua P, Gregorc A, Ivanova E, Kence A, Kence M, Kezic N, Kiprijanovska H, Kozmus P, Kryger P, Le Conte Y, Lodesani M, Manuel Murilhas A, Siceanu A, Soland G, Uzunov A, Wilde J.. A review of methods for discrimination of honey bee populations as applied to European beekeeping. Journal of Apicultural Research. 2011;50(1):51–84.

14. Tarpy DR, Seeley TD. Lower disease infections in honeybee (Apis mellifera) colonies headed by polyandrous vs monandrous queens. Die Naturwissenschaften. 2006;93(4):195–9.

15. Fries I. Nosema ceranae in European honey bees (Apis mellifera). Journal of invertebrate pathology. 2010;103 Suppl 1:S73–9.

16. Hassanein MH. The Influence of Infection with Nosema-Apis on the Activities and Longevity of the Worker Honeybee. Ann Appl Biol. 1953;40(2):418–23.

17. Rinderer TE, Sylvester HA. Variation in Response to Nosema-Apis, Longevity, and Hoarding Behavior in a Free-Mating Population of Honey Bee. Ann Entomol Soc Am. 1978;71(3):372–4.

18. Malone LA, Giacon HA, Newton MR. Comparison of the responses of some New Zealand and Australian honey bees (Apis mellifera L) to Nosema apis Z. Apidologie. 1995;26(6):495–502.

19. Anderson DL, Giacon H. Reduced Pollen Collection by Honey-Bee (Hymenoptera, Apidae) Colonies Infected with Nosema-Apis and Sacbrood Virus. J Econ Entomol. 1992;85(1):47–51.

20. Fries I, Ekbohm G, Villumstad E. Nosema-Apis, Sampling Techniques and Honey Yield. Journal of Apicultural Research. 1984;23(2):102–5.

21. Goodwin M TH A, Perry J, Blackman, R. Cost benefit analysis of using fumagillin to treat Nosema. New Zeal Beekeeper 1990;208:11–2.

22. Genersch E. American Foulbrood in honeybees and its causative agent, Paenibacillus larvae. Journal of invertebrate pathology. 2010;103 Suppl 1:S10–9.

23. Forsgren E. European foulbrood in honey bees. Journal of invertebrate pathology. 2010;103 Suppl 1:S5–9.

24. Ahn AJ, Ahn KS, Noh JH, Kim YH, Yoo MS, Kang SW, et al. Molecular Prevalence of Acarapis Mite Infestations in Honey Bees in Korea. The Korean journal of parasitology. 2015;53(3):315–20.

25. Rosenkranz P, Aumeier P, Ziegelmann B. Biology and control of Varroa destructor. Journal of invertebrate pathology. 2010;103:S96–S119.

26. Mordecai GJ, Wilfert L, Martin SJ, Jones IM, Schroeder DC. Diversity in a honey bee pathogen: first report of a third master variant of the Deformed Wing Virus quasispecies. The ISME journal. 2016;10(5):1264–73.

27. de Miranda JR, Cordoni G, Budge G. The Acute bee paralysis virus-Kashmir bee virus-Israeli acute paralysis virus complex. Journal of invertebrate pathology. 2010;103 Suppl 1:S30–47.

28. Boecking O, Genersch E. Varroosis - the ongoing crisis in bee keeping. J Verbrauch Lebensm. 2008;3(2):221–8.

29. Mondet F, de Miranda JR, Kretzschmar A, Le Conte Y, Mercer AR. On the front line: quantitative virus dynamics in honeybee (Apis mellifera L.) colonies along a new expansion front of the parasite Varroa destructor. PLoS pathogens. 2014;10(8):e1004323.

30. Lively CM, de Roode JC, Duffy MA, Graham AL, Koskella B. Interesting open questions in disease ecology and evolution. The American naturalist. 2014;184 Suppl 1: S1–8.

31. Zheng H, Powell JE, Steele MI, Dietrich C, Moran NA. Honeybee gut microbiota promotes host weight gain via bacterial metabolism and hormonal signaling. Proceedings of the National Academy of Sciences of the United States of America. 2017;114(18):4775–80.

32. Moran NA, Hansen AK, Powell JE, Sabree ZL. Distinctive Gut Microbiota of Honey Bees Assessed Using Deep Sampling from Individual Worker Bees. Plos One. 2012;7(4).

33. Jeyaprakash A, Hoy MA, Allsopp MH. Bacterial diversity in worker adults of Apis mellifera capensis and Apis mellifera scutellata (Insecta: Hymenoptera) assessed using 16S rRNA sequences. Journal of invertebrate pathology. 2003;84(2):96–103.

34. Babendreier D, Joller D, Romeis J, Bigler F, Widmer F. Bacterial community structures in honeybee intestines and their response to two insecticidal proteins. Fems Microbiol Ecol. 2007;59(3):600–10.

35. Martinson VG, Danforth BN, Minckley RL, Rueppell O, Tingek S, Moran NA. A simple and distinctive microbiota associated with honey bees and bumble bees. Mol Ecol. 2011;20(3):619–28.

36. Sabree ZL, Hansen AK, Moran NA. Independent Studies Using Deep Sequencing Resolve the Same Set of Core Bacterial Species Dominating Gut Communities of Honey Bees. Plos One. 2012;7(7).

37. Corby-Harris V, Maes P, Anderson KE. The Bacterial Communities Associated with Honey Bee (Apis mellifera) Foragers. Plos One. 2014;9(4).

38. Engel P, Martinson VG, Moran NA. Functional diversity within the simple gut microbiota of the honey bee. Proceedings of the National Academy of Sciences of the United States of America. 2012;109(27):11002–7.

39. Scardovi VT LD. New species of bifid bacteria from Apis mellifica L. and Apis indica F. A contribution to the taxonomy and biochemistry of the genus Bifidobacterium. Zentralbl Bakteriol Parasitenkd Infektionskr Hyg. 1969;123:64–8.

40. Bottacini F, Milani C, Turroni F, Sanchez B, Foroni E, Duranti S, et al. Bifidobacterium asteroides PRL2011 genome analysis reveals clues for colonization of the insect gut. Plos One. 2012;7(9):e44229.

41. Engel P, Kwong WK, Moran NA. Frischella perrara gen. nov., sp nov., a gammaproteobacterium isolated from the gut of the honeybee, Apis mellifera. Int J Syst Evol Micr. 2013;63:3646–51.

42. Kesnerova L, Moritz R, Engel P. Bartonella apis sp. nov., a honey bee gut symbiont of the class Alphaproteobacteria. Int J Syst Evol Microbiol. 2016;66(1):414–21.

43. Engel P, Moran NA. Functional and evolutionary insights into the simple yet specific gut microbiota of the honey bee from metagenomic analysis. Gut microbes. 2013;4(1):60–5.

44. Engel P, Martinson VG, Moran NA. Functional diversity within the simple gut microbiota of the honey bee. Proc Natl Acad Sci U S A. 2012;109(27):11002–7.

45. Kwong WK, Engel P, Koch H, Moran NA. Genomics and host specialization of honey bee and bumble bee gut symbionts. Proceedings of the National Academy of Sciences of the United States of America. 2014;111(31):11509–14.

46. Lee FJ, Rusch DB, Stewart FJ, Mattila HR, Newton IL. Saccharide breakdown and fermentation by the honey bee gut microbiome. Environmental microbiology. 2015; 17(3):796–815.

47. Forsgren E, Olofsson TC, Vasquez A, Fries I. Novel lactic acid bacteria inhibiting Paenibacillus larvae in honey bee larvae. Apidologie. 2010;41(1):99–108.

48. Engel P, Bartlett KD, Moran NA. The Bacterium Frischella perrara Causes Scab Formation in the Gut of its Honeybee Host. mBio. 2015;6(3):e00193–15.

49. Schmidt K, Engel P. Probiotic Treatment with a Gut Symbiont Leads to Parasite Susceptibility in Honey Bees. Trends in parasitology. 2016;32(12):914–6.

50. Katsnelson A. Microbiome: The puzzle in a bee’s gut. Nature. 2015;521(7552):S56.

51. Aken BL, Achuthan P, Akanni W, Amode MR, Bernsdorff F, Bhai J, et al. Ensembl 2017. Nucleic acids research. 2017;45(D1):D635–D42.

52. Laetsch D.R. BML. Interrogation of genome assemblies [version 1; referees: 2 approved with reservations]. F1000Research. 2017;6:1287.

53. Evans JDS, Ryan; Childers, Anna. HoloBee Database v2016.1. Ag Data Commons2016.

54. Martinez J, Longdon B, Bauer S, Chan YS, Miller WJ, Bourtzis K, et al. Symbionts commonly provide broad spectrum resistance to viruses in insects: a comparative analysis of Wolbachia strains. PLoS pathogens. 2014;10(9):e1004369.

55. Engel P, Kwong WK, McFrederick Q, Anderson KE, Barribeau SM, Chandler JA, et al. The Bee Microbiome: Impact on Bee Health and Model for Evolution and Ecology of Host-Microbe Interactions. mBio. 2016;7(2):e02164–15.

56. Enright AJ, Van Dongen S, Ouzounis CA. An efficient algorithm for large-scale detection of protein families. Nucleic acids research. 2002;30(7):1575–84.

57. Heath LAF. Chalk Brood Pathogens: A Review. Bee World. 1982;63(3):130–5.

58. Ellegaard KM, Tamarit D, Javelind E, Olofsson TC, Andersson SG, Vasquez A. Extensive intra-phylotype diversity in lactobacilli and bifidobacteria from the honeybee gut. BMC genomics. 2015;16:284.

59. Khaled JM, Al-Mekhlafi FA, Mothana RA, Alharbi NS, Alzaharni KE, Sharafaddin AH, et al. Brevibacillus laterosporus isolated from the digestive tract of honeybees has high antimicrobial activity and promotes growth and productivity of honeybee’s colonies. Environmental science and pollution research international. 2017.

60. Plischuk S, Meeus I, Smagghe G, Lange CE. Apicystis bombi (Apicomplexa: Neogregarinorida) parasitizing Apis mellifera and Bombus terrestris (Hymenoptera: Apidae) in Argentina. Environmental microbiology reports. 2011;3(5):565–8.

61. Dias G, Dallai R, Carapelli A, Almeida JP, Campos LA, Faroni LR, et al. First record of gregarines (Apicomplexa) in seminal vesicle of insect. Scientific reports. 2017;7(1):175.

62. Jones JC, Fruciano C, Hildebrand F, Al Toufalilia H, Balfour NJ, Bork P, et al. Gut microbiota composition is associated with environmental landscape in honey bees. Ecology and evolution. 2018;8(1):441–51.

63. Rothschild D, Weissbrod O, Barkan E, Kurilshikov A, Korem T, Zeevi D, et al. Environment dominates over host genetics in shaping human gut microbiota. Nature. 2018.

64. Power SE, O’Toole PW, Stanton C, Ross RP, Fitzgerald GF. Intestinal microbiota, diet and health. The British journal of nutrition. 2014;111(3):387–402.

65. Hamdi C, Balloi A, Essanaa J, Crotti E, Gonella E, Raddadi N, et al. Gut microbiome dysbiosis and honeybee health. J Appl Entomol. 2011;135(7):524–33.

66. Anderson KE, Ricigliano VA. Honey bee gut dysbiosis: a novel context of disease ecology. Current opinion in insect science. 2017;22:125–32.

67. Horton MA, Oliver R, Newton IL. No apparent correlation between honey bee forager gut microbiota and honey production. Peerj. 2015;3.

68. Maes PW, Rodrigues PA, Oliver R, Mott BM, Anderson KE. Diet-related gut bacterial dysbiosis correlates with impaired development, increased mortality and Nosema disease in the honeybee (Apis mellifera). Mol Ecol. 2016;25(21):5439–50.

69. Morimoto T, Kojima Y, Toki T, Komeda Y, Yoshiyama M, Kimura K, et al. The habitat disruption induces immune-suppression and oxidative stress in honey bees. Ecology and evolution. 2011;1(2):201–17.

70. Kwong WK, Mancenido AL, Moran NA. Immune system stimulation by the native gut microbiota of honey bees. Royal Society open science. 2017;4(2): 170003.

71. Gauthier L, Cornman S, Hartmann U, Cousserans F, Evans JD, de Miranda JR, et al. The Apis mellifera Filamentous Virus Genome. Viruses. 2015;7(7):3798–815.

72. Engel P, Stepanauskas R, Moran NA. Hidden diversity in honey bee gut symbionts detected by single-cell genomics. PLoS genetics. 2014;10(9):e1004596.

73. Moran NA. Genomics of the honey bee microbiome. Current opinion in insect science. 2015;10:22–8.

74. Vejnovic B, Stevanovic J, Schwarz RS, Aleksic N, Mirilovic M, Jovanovic NM, et al. Quantitative PCR assessment of Lotmaria passim in Apis mellifera colonies co-infected naturally with Nosema ceranae. Journal of invertebrate pathology. 2018;151:76–81.

75. Bonilla-Rosso G, Engel P. Functional roles and metabolic niches in the honey bee gut microbiota. Current opinion in microbiology. 2018;43:69–76.

76. Graystock P, Meeus I, Smagghe G, Goulson D, Hughes WO. The effects of single and mixed infections of Apicystis bombi and deformed wing virus in Bombus terrestris. Parasitology. 2016;143(3):358–65.

77. Stevanovic J, Schwarz RS, Vejnovic B, Evans JD, Irwin RE, Glavinic U, et al. Species-specific diagnostics of Apis mellifera trypanosomatids: A nine-year survey (2007-2015) for trypanosomatids and microsporidians in Serbian honey bees. Journal of invertebrate pathology. 2016;139:6–11.

78. Bolger AM, Lohse M, Usadel B. Trimmomatic: a flexible trimmer for Illumina sequence data. Bioinformatics. 2014;30(15):2114–20.

79. Li H, Durbin R. Fast and accurate short read alignment with Burrows-Wheeler transform. Bioinformatics. 2009;25(14):1754–60.

80. Li H, Handsaker B, Wysoker A, Fennell T, Ruan J, Homer N, et al. The Sequence Alignment/Map format and SAMtools. Bioinformatics. 2009;25(16):2078–9.

81. DePristo MA, Banks E, Poplin R, Garimella KV, Maguire JR, Hartl C, et al. A framework for variation discovery and genotyping using next-generation DNA sequencing data. Nature genetics. 2011;43(5):491–8.

82. Van der Auwera GA, Carneiro MO, Hartl C, Poplin R, Del Angel G, Levy-Moonshine A, et al. From FastQ data to high confidence variant calls: the Genome Analysis Toolkit best practices pipeline. Current protocols in bioinformatics. 2013;43:11 0 1–33.

83. Sherry ST, Ward MH, Kholodov M, Baker J, Phan L, Smigielski EM, et al. dbSNP: the NCBI database of genetic variation. Nucleic acids research. 2001;29(1):308–11.

84. Cingolani P, Platts A, Wang le L, Coon M, Nguyen T, Wang L, et al. A program for annotating and predicting the effects of single nucleotide polymorphisms, SnpEff: SNPs in the genome of Drosophila melanogaster strain w1118; iso-2; iso-3. Fly. 2012;6(2):80–92.

85. Zheng X, Levine D, Shen J, Gogarten SM, Laurie C, Weir BS. A high-performance computing toolset for relatedness and principal component analysis of SNP data. Bioinformatics. 2012;28(24):3326–8.

86. Bankevich A, Nurk S, Antipov D, Gurevich AA, Dvorkin M, Kulikov AS, et al. SPAdes: a new genome assembly algorithm and its applications to single-cell sequencing. Journal of computational biology: a journal of computational molecular cell biology. 2012;19(5):455–77.

87. Morgulis A, Coulouris G, Raytselis Y, Madden TL, Agarwala R, Schaffer AA. Database indexing for production MegaBLAST searches. Bioinformatics. 2008;24(16):1757–64.

88. Coordinators NR. Database resources of the National Center for Biotechnology Information. Nucleic acids research. 2018;46(D1):D8–D13.

89. UniProt Consortium T. UniProt: the universal protein knowledgebase. Nucleic acids research. 2018;46(5):2699.

90. Buchfink B, Xie C, Huson DH. Fast and sensitive protein alignment using DIAMOND. Nature methods. 2015;12(1):59–60.

91. Theocharidis A, van Dongen S, Enright AJ, Freeman TC. Network visualization and analysis of gene expression data using BioLayout Express(3D). Nature protocols. 2009;4(10):1535–50.

92. Purcell S, Neale B, Todd-Brown K, Thomas L, Ferreira MA, Bender D, et al. PLINK: a tool set for whole-genome association and population-based linkage analyses. American journal of human genetics. 2007;81(3):559–75.

93. Alexander DH, Novembre J, Lange K. Fast model-based estimation of ancestry in unrelated individuals. Genome research. 2009;19(9):1655–64.

94. Harpur BA, Kent CF, Molodtsova D, Lebon JM, Alqarni AS, Owayss AA, et al. Population genomics of the honey bee reveals strong signatures of positive selection on worker traits. Proceedings of the National Academy of Sciences of the United States of America. 2014;111(7):2614–9.

95. Neuditschko M, Khatkar MS, Raadsma HW. NetView: a high-definition network-visualization approach to detect fine-scale population structures from genome-wide patterns of variation. Plos One. 2012;7(10):e48375.

96. Chen Y, Evans JD, Smith IB, Pettis JS. Nosema ceranae is a long-present and wide-spread microsporidian infection of the European honey bee (Apis mellifera) in the United States. Journal of invertebrate pathology. 2008;97(2):186–8.

97. Dixon P. VEGAN, a package of R functions for community ecology. J Veg Sci. 2003;14(6):927–30.

98. Quast C, Pruesse E, Yilmaz P, Gerken J, Schweer T, Yarza P, et al. The SILVA ribosomal RNA gene database project: improved data processing and web-based tools. Nucleic acids research. 2013;41(D1):D590–D6.

